# Sex differences and behaviour in the pace-of-life of rodents

**DOI:** 10.1101/2024.08.15.608175

**Authors:** Bryan Hughes, Gabriela F. Mastromonaco, Jeff Bowman, Albrecht Schulte-Hostedde

## Abstract

Male and female rodents experience different selective pressures associated with reproductive costs. Thus, we may expect the expression of different Pace-of-life (POL) strategies between sexes. Further, the pace-of-life syndrome (POLS) hypothesis and anisogamy predict differences in the costs of gamete production, where variation in life history trait expression should follow a fast-slow continuum such that males and females might exist on opposite ends of the spectrum. However, males and females could express a similar POL strategy in systems where the reproductive costs are consistent between sexes or where selective pressures force a similar directionality of traits. We used standardized behavioural tests and fecal glucocorticoid metabolite (FGM) concentrations to measure potential differences in POL strategies among three rodent species in Algonquin Provincial Park. We hypothesized that differences in reproductive costs between males and females would result in differences in POL traits along the fast-slow continuum. We predicted that males would express more explorative behaviours and have a higher level FGMs than female counterparts. We found little support for the POLS hypothesis between sexes. Instead, we suggest that the species observed may express a similar directionality of selection for the observed behavioural traits, where both sexes express similar relationships in POL. Thus, some male rodents may be more explorative to accommodate the increase of energetic stress associated with mate acquisition, while females may share similar trait expressions to accommodate the increased metabolic demand for the care and development of young.

**Significance statement:** Docility and exploration are measurements of how species or individuals react to novel and sometimes risky stimuli. Further, reproduction is an energetically expensive and inherently risky process. In seasonally breeding rodents, males often invest in fast pace-of-life behaviours that accommodate an increased necessity to disperse and find a mate. Meanwhile, females often invest in metabolically expensive physiological processes, including pregnancy and lactate development, increased protection of the young and the need for self-maintenance. Using a series of behavioural tests and fecal glucocorticoid metabolites (FGMs), we measured the relationship between metabolic stress, docility and exploration in three species of rodents during the breeding season. We observed consistent relationships in behaviour and FGMs, indicating uniformity between sexes regarding these behaviours. We posit that the costs associated with mate acquisition for males and the care and development of young for females can influence similar behavioural strategies.

## Introduction

The pace-of-life syndrome (POLS) hypothesis argues that trade-offs in energetic investment stemming from resource allocation should result in differences in behavioural, physiological, and life-history strategies across species. Relationships among POLS traits are predicted to evolve along a fast-slow continuum (Réale et al. 2010). Fast-paced species are predicted to have a high reproductive output with low investment in parental care, express explorative and bold behaviours, and invest less in self-regulation, including a lower metabolic reactivity (Réale et al. 2010; Santostefano et al. 2017; Hall et al. 2015). Slow-paced species display traits that reflect the opposite, with a lower reproductive output yet higher investment in individual young (Réale et al. 2010). The POLS hypothesis may also be used to evaluate individual differences in behavioural strategy within species (Dammhahn et al. 2018). Males and females often express different reproductive roles within a population resulting in sex-specific variation and energetic trade-offs regarding self-maintenance and reproduction (Hämäläinen et al. 2018; Tarka et al. 2018). Differences in energetic trade-offs drive natural and sexual selection, resulting in sexually dimorphic traits or sexual differences in behaviour and physiology that might influence the position of individuals along the predicted fast-slow continuum (Shutler 2010; Hedrick and Temeles 1989; Fairbairn et al. 2007). Indeed, there is evidence of variation in life-history traits between sexes, including size, age of maturation, growth, reproductive potential, and mortality (Martin 2004; Clutton-Brock and Vincent 1991; Bonduriansky et al. 2008; Maklakov and Lummaa 2013; Adler and Bonduriansky 2014; Arso Civil et al. 2019). Although many taxa have sex-specific life-history traits, the directionality and expression of sex-specific traits and syndromes remains less understood (Tarka et al. 2017).

Trait expression between sexes may stem from sex-specific reproductive roles influenced by anisogamy (Lehtonen et al. 2016). Anisogamy suggests that behavioural and physiological traits in males support a greater investment in mate acquisition, while females invest more in self-maintenance and the development of offspring (Schärer et al. 2012; Lehtonen et al. 2016). Thus, males often occupy the fast end of the POLS hypothesis compared to female counterparts (Tarka et al. 2018). Sex-specific variation has been proposed to be one of the underlying mechanisms responsible for a lack of empirical support for the assumptions presented by the POLS hypothesis within taxa (Royauté et al. 2018; Hämäläinen et al. 2018; Immonen et al. 2018). Given the association between POL traits, sex can be an important variable when evaluating various physiological or behavioural phenotypes. Selective pressures should result in alternative physiological patterns, including immunity (Monceau et al. 2017; Love et al. 2008; Lee 2006), metabolism (Shingleton and Vea 2023; Rønning et al. 2016), hormone production (Nelson, 2005), and thermoregulation (van Jaarsveld et al. 2021) between sexes. Consequently, selective pressures should also favour alternative behavioural strategies, including in risk-taking behaviours (Holtby and Healey 1990), foraging patterns (De Pascalis et al. 2020) and aggression (Fresneau et al. 2014). Given the assumptions of anisogamy, males should express behavioural and physiological traits differently than females, including higher exploration and risk-taking and reduced physiological reactivity to environmental stimuli.

The POL syndrome predicts a correlation between physiological mechanisms and behavioural strategies (Réale et al. 2010). In ecology, stress is the energetic change and physiological response to environmental stimuli (Romero, 2004; Hobfoll 1988; Costantini 2008). Fecal glucocorticoid metabolites (FGMs) are a standard non-invasive proxy of metabolic stress in animals (Palme 2019; Palme 2012; but see MacDougall-Shackleton et al. 2019 for an overview of why glucocorticoids are not “stress” hormones specifically). Glucocorticoids, such as corticosterone and cortisol, are a group of steroid hormones released from the hypothalamic-pituitary-adrenal axis (HPA-axis) that assist in regulating metabolic function in response to external stimulus (Palme 2019; Toufexis et al. 2014). Indeed, a fast POLS is often associated with increased exploration and risk-taking behaviour (more reactive strategies) to accommodate mate acquisition (Réale et al. 2010). Consequently, increased exploration and risk-taking behaviour benefits from a lower investment in energetic reactivity to environmental stimuli – thus fast paced individuals express a lower fluctuation of FGMs during novel events (Boyce and Ellis 2005). Between sexes, this may suggest that males should be more exploratory and less docile compared to female conspecifics. In contrast, the slow end of the POLS often invest in strategies that prioritize self-preservation (more proactive behaviours), including an increased reactivity to environmental stimuli. Indeed, a greater investment in physiological functions relating to offspring development (pregnancy and lactate production) are metabolically expensive and require increased energetic investment during breeding (Reeder and Kramer 2005). Thus, between sexes males are predicted to express more reactive strategies, compared to more proactive female counterparts.

Hämäläinen et al. (2018) provide a framework that suggests sex-specific POLS traits may evolve from various circumstances where predicted covariance may be expressed differently between sexes. Anisogamy has been well described in rodents (Ramm et al. 2005; Dewsbury, 1982; Roldan et al. 1992), where males and females often express alternative coping strategy and physiology based on energetic demands during reproduction (Eccard and Herde 2013; Immonen et al. 2018). We quantified sexual differences in movement behaviour and energetic investment, measured through FGMs (Palme et al. 2019), between sexes in three species of rodent, including the deer mouse (*Peromyscus maniculatus*), the red-backed vole (*Clethrionomys gapperi*) and the woodland jumping mouse (*Napaeozapus insignis*). We hypothesized that reproductive role should lead to alternative expression of traits between sexes according to the POLS. If our hypothesis is supported, we predicted that males should express more reactive behavioural phenotypes, including being more exploratory and less docile than females within the same species. Understanding sex-specific variation in POL traits is one step in answering broader questions concerning the evolution of temperament and how competition between conspecifics is mitigated given different selective pressures within the environment.

## Methods

### Capture and handling of animals

Deer mice, red-backed voles, and woodland jumping mice were surveyed in Algonquin Provincial Park, Ontario, Canada, from May through September 2022 across 17 traplines following animal care protocols (#6011106) approved by Laurentian University. Each trapline consisted of 20 Sherman traps (H. B. Sherman Traps, Inc., Tallahassee, Florida) baited with water-soaked sunflower seeds. Traplines were composed of 100-m transects with two traps placed every 10 m, covering an array of forest habitats (see Fryxell et al. 1998 for detailed trapline methodology and habitat description). Traps were baited at dusk and checked the following morning, and each trapline was baited biweekly for 3 consecutive nights. Individuals received metal ear tags with unique alphanumeric codes (National Band and Tag Co., Newport, Kentucky, USA) for identification. For each individual, we recorded the sex (male or female), age class (juvenile, sub-adult, or adult) based on body mass and hair colour (Schmidt et al. 2019), body mass using a 0.1 g Pesola scale, and reproductive status measured as scrotal or non-scrotal (absence or presence of testes) for males, and non-reproductive, pregnant, perforate or lactating (no visible signs of reproduction, enlarged abdomen, in estrous, or enlarged mammary glands) for females.

### Behavioural tests

Individuals were introduced to one behavioural test during a day. Each test was used to quantify a single behavioural temperament, including docility (an individual’s likelihood to engage with perceived risks from predators) from the Handling Bag Test (Martin and Réale 2008) or exploration and activity (an individual’s likelihood to engage in risky non-novel stimuli) from the Open Field Test (Carter et al., 2013). We recorded each test using a camera (Sony HDR-CX405) and later analyzed behaviours using CowLog 3.0 (Pastell 2016; see supplementary data A and B for ethograms). Due to all behavioural tests occurring in the wild, data was not recorded in a blind manner. However, behavioural videos were quantified at a later date by the same observer to standardize the analyses of each test.

### Fecal collection, extraction, and immunoassay analyses

All materials used for the immunoassays were purchased from Avantor Sciences unless otherwise specified. Fecal samples were used as a non-invasive measure of glucocorticoid levels in all three species. All materials used for the collection of fecal samples were purchased from Fisher Scientific. Small mammals often defecate in response to handling, therefore fecal samples were collected immediately after defecation during the handling process and placed in an Eppendorf tube filled with 80% methanol. Eppendorf tubes were then placed on ice in a cooler and were later transferred to a -20 °C freezer for temporary storage during our four-month collection period. All samples were later transferred to a -80 °C in September, where they were stored until immunoassay analyses were performed. Where not possible to collect fecal samples during handling due to insufficient sample volume or lack of defecation, fecal samples were collected from the Sherman trap, no later than 19 h after defecation (Veitch et al. 2021). While traplines were checked in the same order each week to reduce decay of fecal samples, the 19-hour collection period has no significant impact on total FGM concentrations (Veitch et al. 2021). Species specific differences in metabolic activity and fecal glucocorticoids are observed through radiometabolic assays henceforth referred to as a biological validation (Palme 2019). Although no such validation has previously been performed for woodland jumping mice, enzyme immunoassays for measuring FGMs have been validated for red-backed voles (Eleftheriou et al. 2020) and deer mice (Harper and Austad. 2000).

Glucocorticoid metabolites were extracted using methods detailed by Veitch et al. (2021) with some modifications. Each fecal sample along with any associated methanol from the collection was transferred to a 7 ml glass vial. An additional 1 ml of 100 % methanol was used to rinse out the collection tube and was then added to the same glass vial. Samples were left under a fume hood to completely evaporate the alcohol to obtain a fecal sample mass. Freshly prepared 80 % methanol was added to the samples using a ratio of 0.05 g/ml before vortexing samples for 10 s. Vortexed samples were placed on an orbital shaker at 100 rpm for approximately 20 h. The following day, the vials were centrifuged for 10 min at 2400 g and the supernatants were transferred into fresh glass vials and placed at - 20 °C until analysis.

Enzyme immunoassays (EIAs) were quantified using methods described in Baxter-Gilbert et al. (2014) and Stewart et al. (2020). Species specific differences in fecal hormone metabolite profiles are typically assessed using biochemical, physiological and biological validation techniques that ensure appropriate selection of EIAs (Palme 2019). EIAs for measuring FGMs have been previously validated for red-backed voles (Eleftheriou et al. 2020) and deer mice (Harper and Austad. 2000), while no such validation has previously been performed for woodland jumping mice. However, based on the evolutionary similarity and phylogenetic relationships between woodland jumping mice, deer mice and red-backed voles (i.e. species that are validated), we do not expect significant differences in hormone metabolite excretion. Deer mouse and red-backed vole FGM concentrations were quantified using modifications of a corticosterone enzyme immunoassay (EIA) previously described (Veitch et al. 2021) and woodland jumping mice were quantified using both a corticosterone and cortisol EIA. Extracts were diluted in EIA buffer at 1:30 and 1:20 for deer mouse and red-backed vole, respectively. Antibody and horseradish peroxidase conjugate were diluted 1:300,000 and 1:1,000,000 in EIA buffer, respectively. Absorbance was measured using a spectrophotometer (Epoch 2 microplate reader, BioTek, Winooski, VT, USA). Intra-assay CV was 7.5%, and inter-assay CVs were 6.6% and 5.9% (45% binding) for deer mouse and red-backed vole respectively, and 8.2% and 12.4% (70% binding) for deer mouse and red-backed vole, respectively. Woodland jumping mouse FGMs were quantified using a cortisol enzyme immunoassay (EIA) adapted from Munro and Stabenfeldt (1984) and modified by Edwards et al. (2019) for a double antibody EIA with some additional modifications. Briefly, extracts were diluted in assay buffer (0.1 M Tris buffer, pH 7.5, containing 0.15 M NaCl, 10 mM EDTA, 0.1% bovine serum albumin, and 0.1% Tween 20) at 1:10 to 1:100. An in-house made blocking solution (250 μl; 10 mM phosphate, 15 mM NaCl, 1% sucrose, 2% bovine serum albumin, and 0.01% Tween 20, pH 7.5) was added to the wells and incubated for 4-24 hours at RT, then aspirated. Plates were dried at RT in a desiccator cabinet (Dry Keeper cabinet, SP Bel-Art, Wayne, NJ, USA) until relative humidity (RH) was below 20%. GARG coated plates remained in desiccator cabinet (≤ 20% RH) until immediately before use. Pre-coated GARG plates were loaded with 50 μl cortisol standard (Sigma H0135; 19.6-20,000 pg/ml), diluted extracts and controls, followed by 50 μl horseradish peroxidase conjugate (1:300,000) and 50 μl cortisol antiserum (1:300,000; antibody R4866; C. Munro, University of California, Davis, CA, USA), all diluted in assay buffer. Plates were incubated for 2 hours at RT in the dark, and then washed with 0.05% Tween 20, 7.5 mM NaCl, 0.01 mM EDTA, 5 mM phosphate solution (pH 7.2). To each well, 200 μl of substrate solution (0.5 ml of 4 mg/ml tetramethylbenzidine in dimethylsulphoxide and 0.1 ml of 0.176 M H_2_O_2_ diluted in 22 ml of 0.01 M sodium acetate trihydrate [C_2_H_3_NaO_2_ ·3H_2_O], pH 5.0) was added. After 30 min incubation in the dark, colour reaction was stopped with 50 μl H_2_SO_4_ (1.8 M) and absorbance was measured at 450 nm with a reference of 630 nm spectrophotometrically. Intra-assay CV was 11.0%, and inter-assay CVs were 5.5% (11% binding) and 10.4% (35% binding).

Parallel displacement between the standard curve and serial dilutions of fecal extract was used as an indirect measure of assay specificity and a biochemical validation of the selected EIAs. Pooled reconstituted fecal extracts were serially diluted two-fold in assay buffer and compared to the respective standard curve. The data were plotted as log (relative dose) vs. percent antibody bound (Microsoft Excel). The slopes of the lines within the linear portion of the curves were determined using linear regression analysis and compared (Soper 2021) where p > 0.05 indicates that the slopes are not significantly different and thus interpreted as parallel. For Jumping Mice, serial dilutions of pooled fecal extract showed parallel displacement with the cortisol standard curve (t=0.10, p=0.92, df=5; supplementary data C), but not with the corticosterone standard curve (t=5.97, df = 7, P = <0.0001, supplementary data C). In contrast, serial dilutions of pooled fecal extract showed parallel displacement with the corticosterone standard curve for deer mice (t = 0.71, df = 9, P = 0.50, Supplementary Fig A2), and red-backed voles (t = 0.03, df = 9, P = 0.98, supplementary data C). Samples were assayed at the dilution factor that corresponded to 50% binding of the serially diluted fecal pool for each assay. Cross reactivities for corticosterone antibody (CJM006) and cortisol antibody (R4866) are reported in Metrione & Harder (2011) and Young et al. (2001), respectively. The double antibody EIAs used in jumping mice were used because no previous validation for glucocorticoids have been described in this species. The parallel displacement for the cortisol EIA showed a greater binding specificity and is therefore the reported parallelism in this study. In some instances, EIA specificity may be better at detecting non-dominant glucocorticoids such as the results reported by Carlsson et al. (2016).

### Statistical analyses

All statistical analyses were conducted using the statistical software program R version 4.2.3 (R Core team 2023). A log_10_ transformation was applied to the total FGM concentrations variable for all models to normalize distribution of data. FGMs outside the high and low cut-off values obtained using the Soper (2021) standard curve calculation are considered inaccurate and were thus removed as outliers (deer mice = > 9500 or < 90 ng / g, Woodland-Jumping Mice >2500 or < 20 ng / g, red-backed voles = >6200 or < 60 ng / g). Low cut-offs were determined as the limit of quantitation (LOQ) using the blank determination method described in Shrivastava and Gulpa (2011), high cut-off values were determined through visual assessment of where values exceeded the standard curve of the parallelism (supplementary data C). Individuals that had multiple fecal samples from the same trap week (and thus same reproductive period) were averaged to evaluate FGM concentrations during that period. We then coupled behavioural tests during the same week of collection to analyze the relationship between behaviour and FGMs (deer mice docility n = 104, and exploration n = 45; red-backed vole docility = 52 and exploration = 27, woodland jumping mouse docility = 14 and exploration = 10).

Linear mixed effects models we used with individual ID as a random effect for each species separately using the lme4 package in R (Bates et al 2015). Independent variables representing potential variation in FGMs between males and females of each species were used, including age class (adult, sub-adult or juvenile), reproductive status (scrotal, non-scrotal, pregnant, lactating, or non-reproductive), body mass and date (May through August). Finally, a linear mixed effect model with all predictor variables was used to show the relationship between behaviour and log_10_ transformed FGMs.

## Results

We found a high degree of variation in the FGM concentrations between individuals for all three species (Figure 1). When comparing FGMs among species based on reproductive status we showed that there was no significant differences in FGMs between pregnant, lactating or non-actively reproductive females, or scrotal and non-scrotal males (Figure 2). Similarly, we observed no significant differences in FGMs between non-reproductive females and non-scrotal males in any of the three study species.

**Figure 1:**
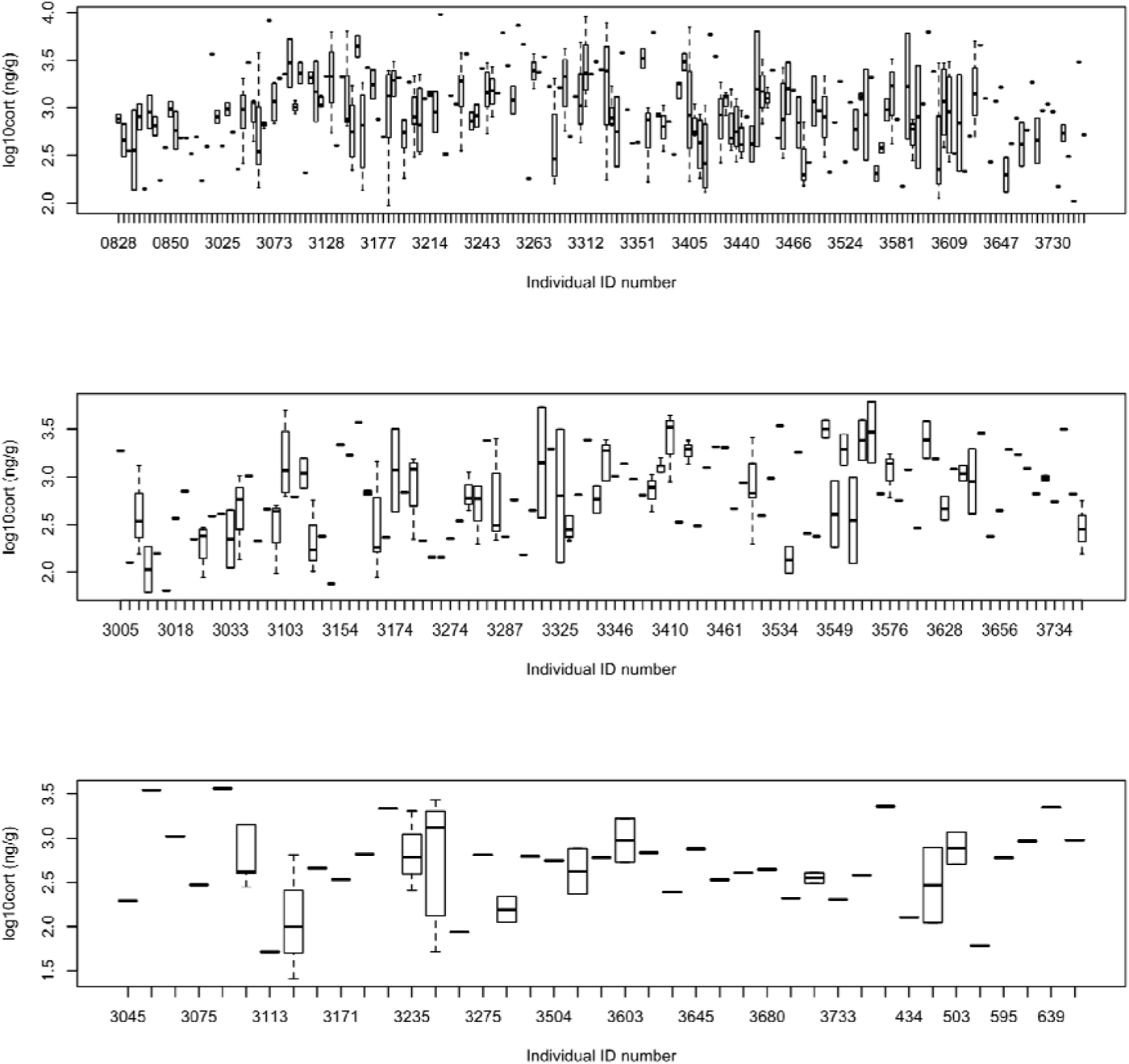
Individual variation in log_10_-transformed FGM in each observed species. Shown are the median (black line) interquartile range (box) and minimum/maximum values (bars). All species show considerable variation in FGMs.

**Figure 2:**
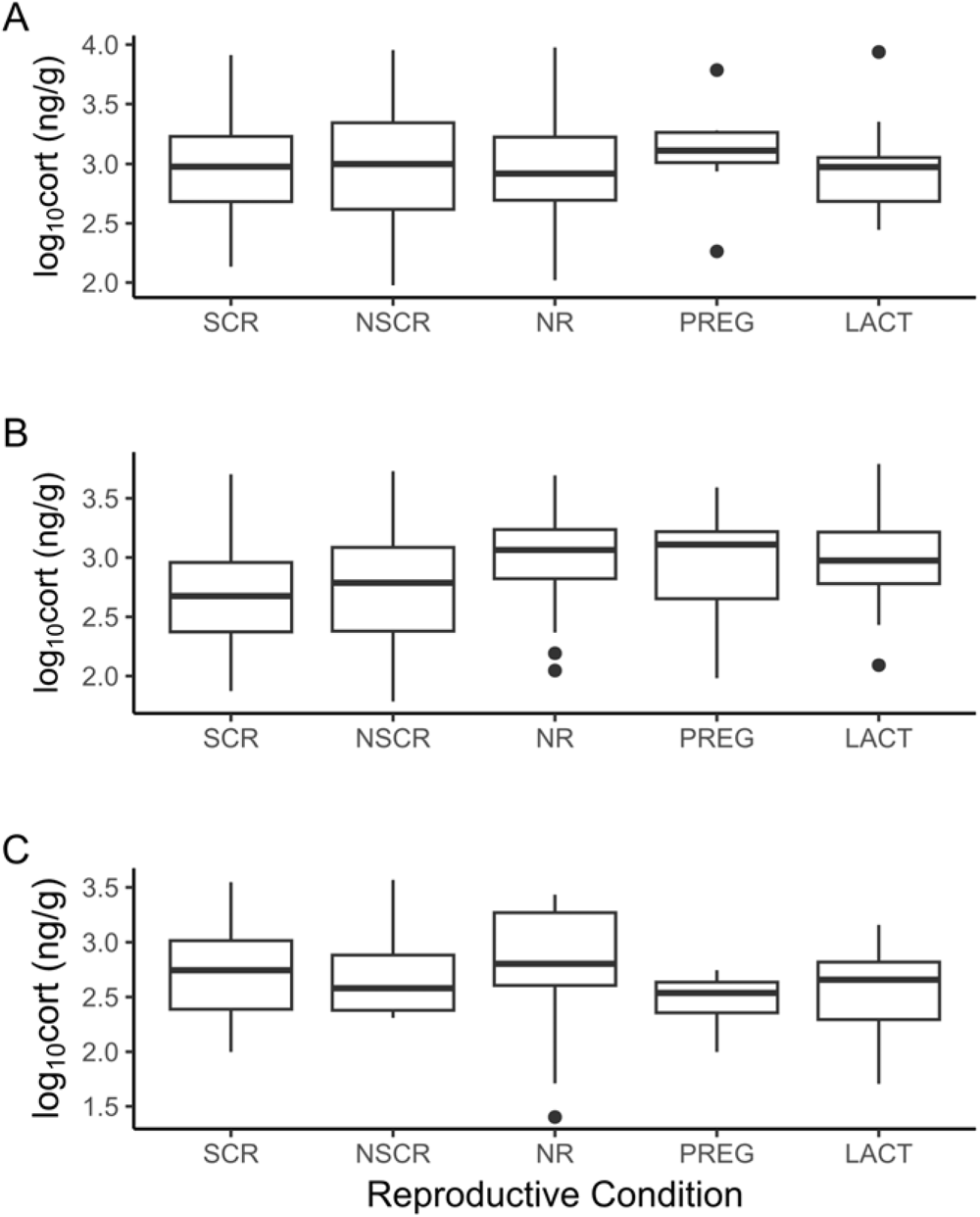
A series of boxplots showing the total basal FGM in deer mice (panel A), red-backed voles (panel B) and woodland jumping mice (panel c). Each column represents reproductive status (NSCR = non-scrotal, SCR = scrotal for males and PREG = pregnant, LACT = lactating, PERF = perforate, or NR = non-reproductive for females). Dark lines represent the mean value of FGMs (ng/g) while whiskers represent the standard deviation.

### Docility and FGMs

We measured the relationship between total docility (time spent immobile) and log_10_-transformed FGM using linear models for a total of 104 deer mice (F = 1.605, df = 92, P = 0.11, R^2^ = 0.16), 15 woodland jumping mice (F = 0.89, df = 38, P = 0.56, R^2^ = 0.22) and 52 red-backed voles (F = 18.59, df = 4, P = 0.0062, R^2^ = 0.98). Our results showed non-statistically significant values using docility as a dependent variable for most predictor variables (date, reproductive status, body mass, sex, and age class), in deer mice and red-backed voles (Table 1; Figure 3). However, body mass and pregnancy was significantly associated with docility in deer mice (P = 0.045), while reproductive condition (P = 0.0044), date (P = 0.0092) and mass (P = 0.0039) were significantly associated with docility in woodland jumping mice (Table 1).

**Table 1:**
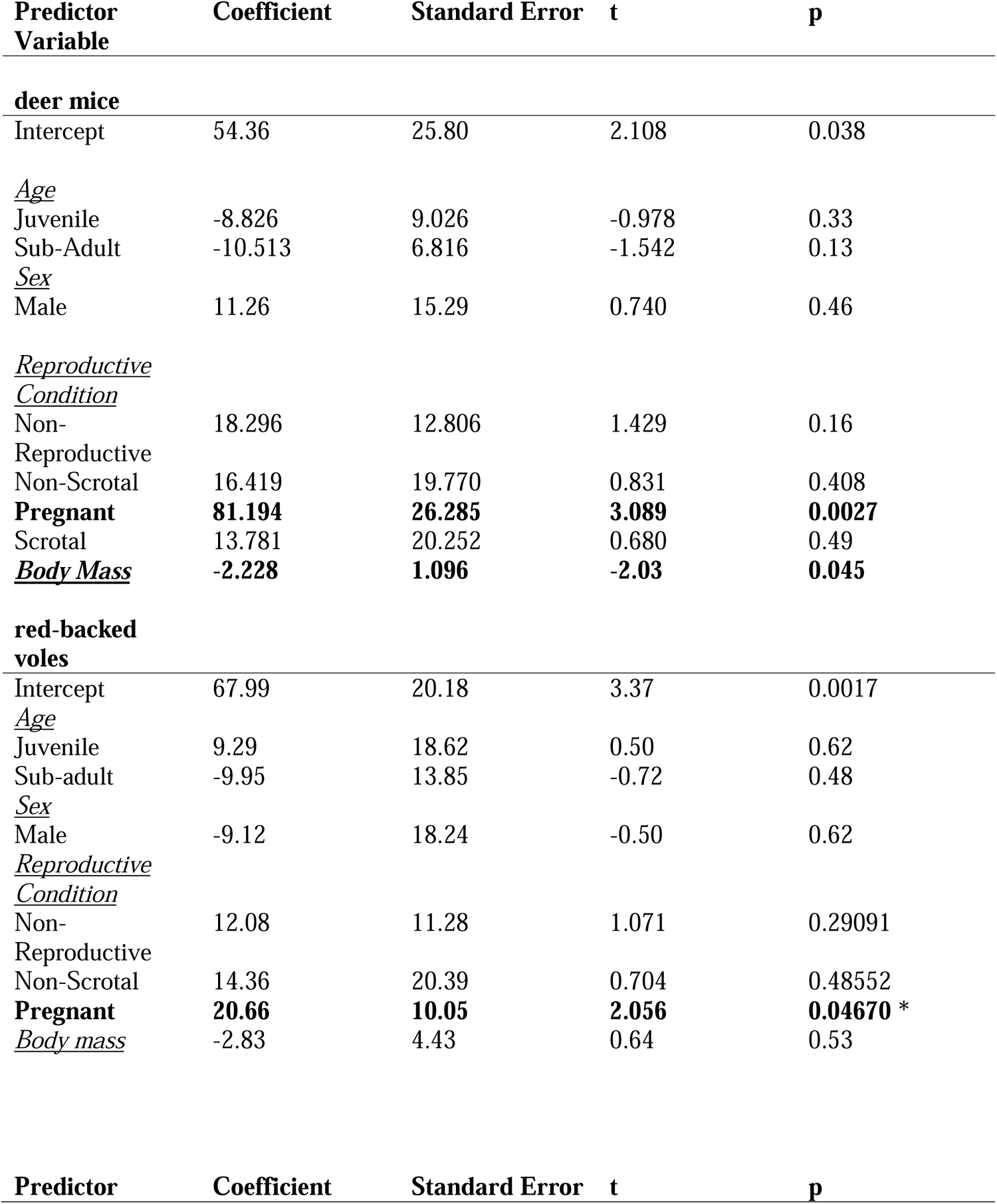
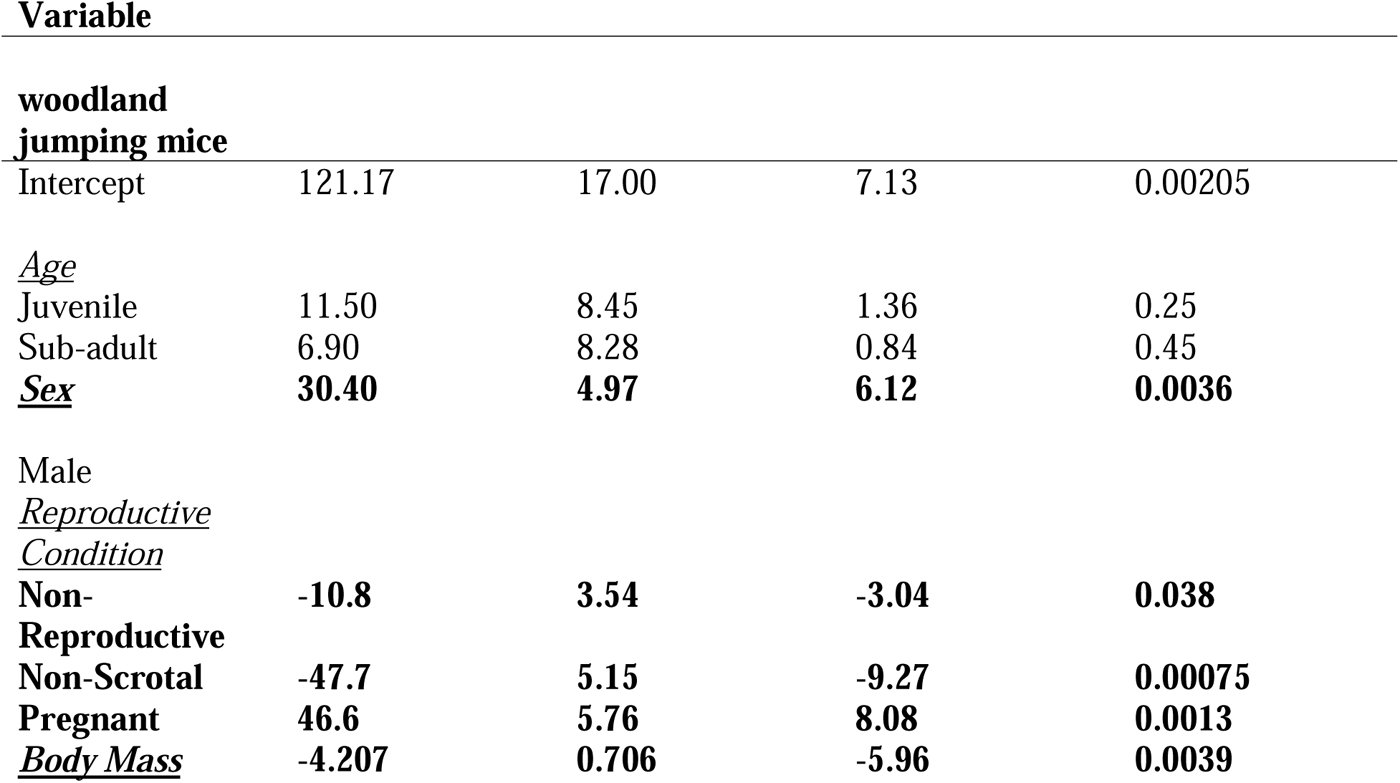
Linear regression statistics for the relationship between docility, measured through the Handling Bag Tests (BT) and log_10_ transformed FGM in deer mice (n = 104), red-backed voles (n=52) and woodland jumping mice (n = 14), using sex, age class (adult, sub-adult and juvenile), reproductive condition, test month and body mass as random effects.

**Figure 3:**
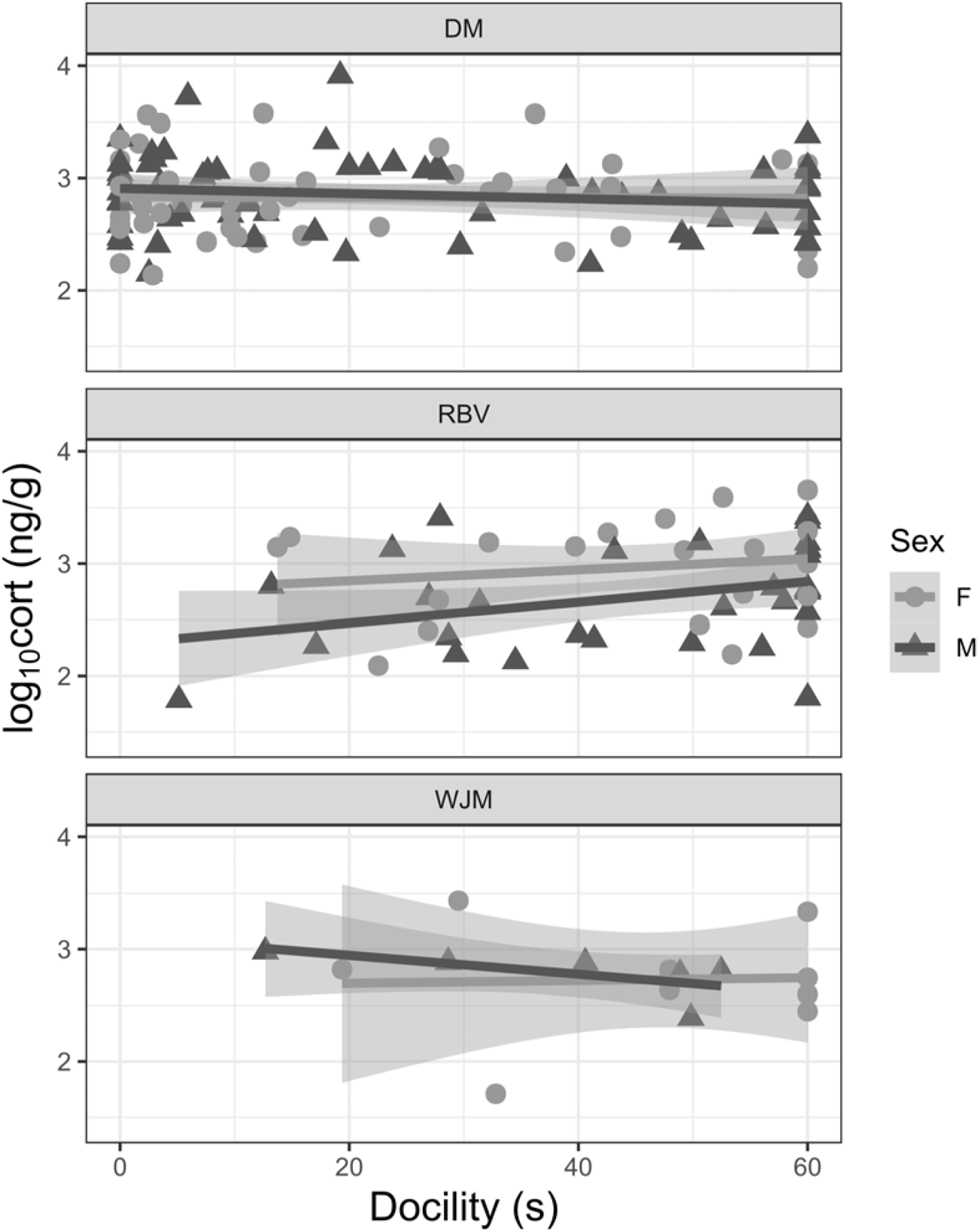
Relationship between log_10_-transformed values of fecal metabolites in deer mice and docility. Individuals showed a negative relationship between docile behaviour and basal log_10_ FGM in deer mice and male Jumping Mice, however red-backed voles showed a positive relationship (total n = 104, males = 61, females = 44) woodland jumping mice (total n = 14, males = 6, females - 8) and red-backed voles (total n = 52, males =31, female = 21). Data points are jittered and represent individual test (red circle for females, and blue triangle for males). 95% confidence intervals are shown by shading.

When comparing within-category effects for the predictor variables (date, reproductive status, body mass, sex, and age class), pregnant individuals had the most significant relationship with total docility (P=0.012; Figure 4); however, this result was most prominent in red-backed voles. Pregnant red-backed voles showed a positive relationship between total FGMs and docility, such that individuals with a higher concentration of FGM expressed more frequent motionless behaviour (Figure 4).

**Figure 4:**
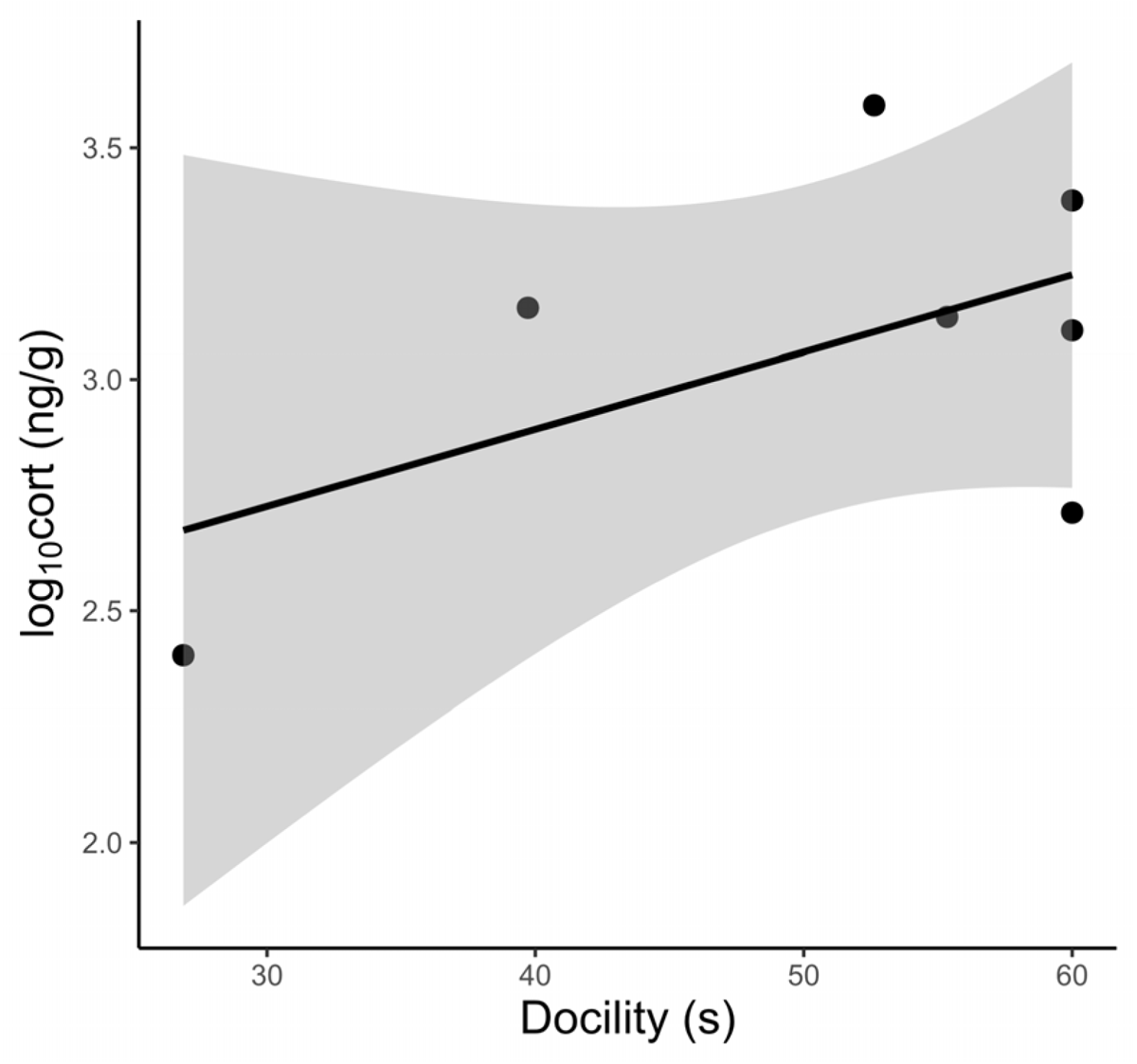
The relations hip between log_10_-transfor med values of FGMs and docility in pregnant red-backed voles. Individual s that underwent a Handling Bag Test are shown via black dots, and 95% confidence interval extracted through linear mixed effects model is represented through shading. red-backed voles show a positive relationship between docility and FGMs, where individuals with a higher concentration of glucocorticoids express higher levels of immobilization, or docile behaviour consistent with the POLS hypothesis.

### Exploration and FGMs

Across species, there were negligible differences in exploration between sexes (Table 3 and Figure 5). We measured the relationship between total exploratory behaviour (time spent moving around an Open Feld Test) and the log_10_ transformed FGMs across a total of 46 deer mice (F = 1.34, df = 35, P = 0.25, R^2^ = 0.276), 10 woodland jumping mice (F = 0.36, df = 3, P = 0.86, R^2^ = 0.36) and 27 red-backed voles (F = 2.654, df = 15, P = 0.042, R^2^ = 0.63). Of the three species, only red-backed voles showed sexual differences in exploration behaviour, such that male red-backed voles were less explorative than female counterparts. Sub-adult deer mice also had a significant relationship with total exploration time (P = 0.0028) such that sub-adult individuals expressed a greater amount of exploration compared to juvenile and adult deer mice and had the greatest concentration of FGMs (Table 3 and Figure 6).

**Figure 5:**
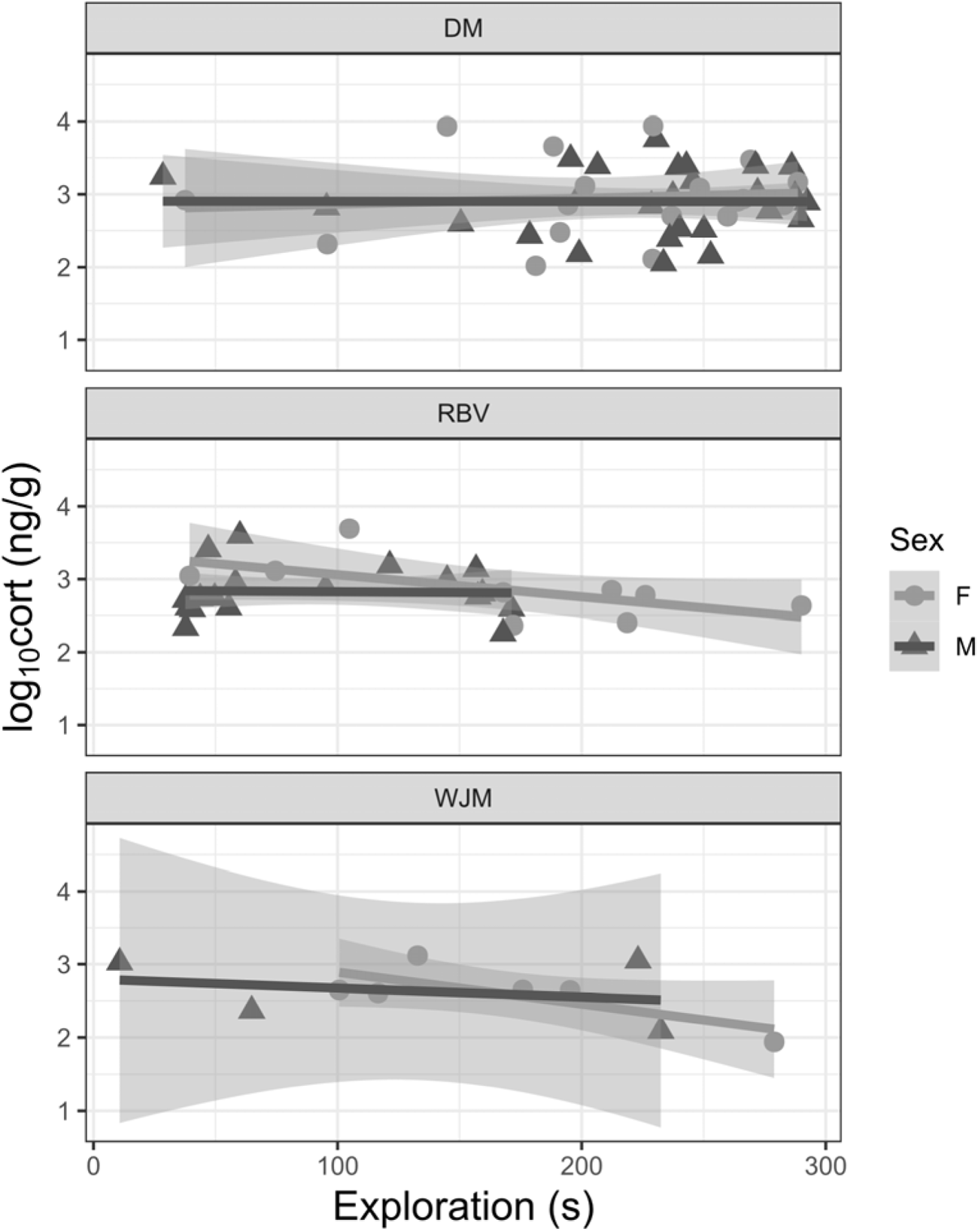
The relationship between log_10_-transformed fecal metabolites and total exploration behaviour in deer mice. Individuals showed a negative relationship in Jumping Mice, Red Backed Voles, and male deer mice; however, a positive relationship in female deer mice (total n = 45, males = 25, females = 19; red-backed voles total n = 27, males = 18, females = 9; woodland jumping mice total n = 10, males = 4, females = 6). Data points are shown by jitters (red circle for females, and blue triangles for males). Shading represents 95% confidence interval.

**Figure 6:**
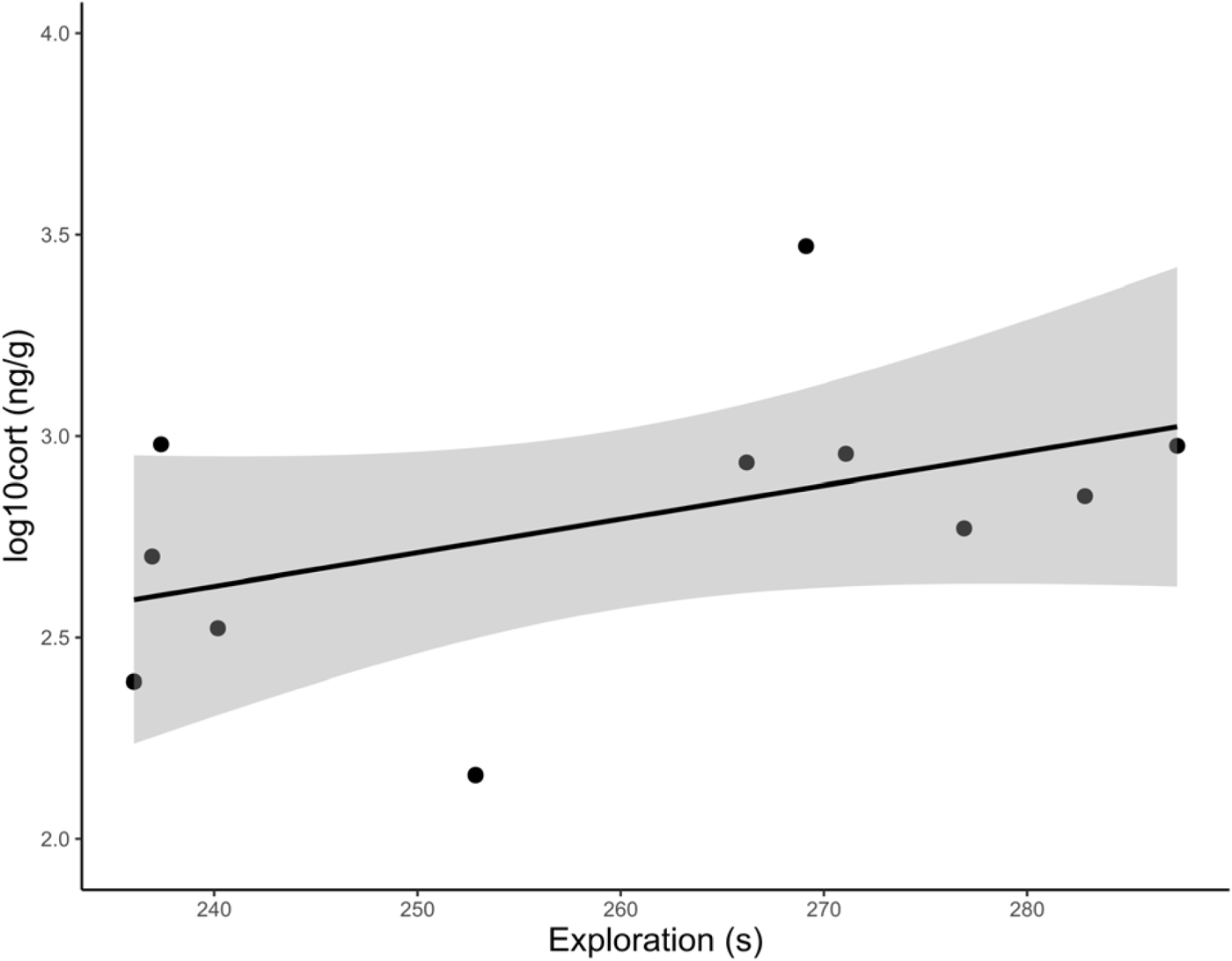
The relationship between log_10_-transformed values of FGMs and exploration in sub-adult deer mice. Exploration in seconds, measured by the Open Field Test are shown via black dots, and 95% confidence interval extracted through linear mixed effects model is represented through shading. Deer mice show a positive relationship between exploration and basal FGMs, where individuals with a higher concentration of glucocorticoids express higher levels of exploration.

**Table 3:** Linear regression statistics for models examining the relationship between total exploration time and log_10_ transformed FGM in deer mice (n = 45), red-backed voles (n = 27) and woodland jumping mice (n = 10) using age (excluded for woodland jumping mice), sex, date and reproductive condition as predictor variables. Significant values (P < 0.05) are shown in bold.

| **Predictor Variable** | **Coefficient** | **Standard Error** | **t** | **P** |
| --- | --- | --- | --- | --- |
| **deer mice** |  |  |  |  |
| Intercept | 182.63 | 93.18 | 1.96 | 0.0582 |
| *Sex* |  |  |  |  |
| Male | -13.24 | 53.578 | -0.25 | 0.80 |
| *Age* |  |  |  |  |
| Juvenile | -8.29 | 30.298 | -0.274 | 0.79 |
| **Sub-Adult** | **42.01** | **26.657** | **1.576** | **0.012** |
| *Reproductive Condition* |  |  |  |  |
| Non-Reproductive | -26.076 | 52.308 | -0.499 | 0.62 |
| Non-Scrotal | -5.244 | 32.82 | -0.160 | 0.87 |
| *Body Mass* | 2.975 | 3.768 | 0.790 | 0.44 |
| **red-backed voles** |  |  |  |  |
| Intercept | 363.90 | 110.8 | 3.28 | 0.0502 |
| *Sex* |  |  |  |  |
| **Male** | **-146.91** | **46.34** | **-3.170** | **0.0063** |
| *Age* |  |  |  |  |
| Sub-Adult | 0.01709 | 59.55 | 0.0001 | 0.99 |
| *Reproductive Condition* |  |  |  |  |
| Non-Reproductive | -102.93 | 57.95 | -1.78 | 0.096 |
| Non-Scrotal | -58.38 | 32.04 | -1.82 | 0.89 |
| Pregnant | -72.38 | 67.96 | -1.07 | 0.30 |
| *Body Mass* | -2.28 | 4.44 | -0.64 | 0.53 |
| woodland jumping mice |  |  |  |  |
| *Sex* |  |  |  |  |
| Male | -194.94 | 360.66 | -0.541 | 0.626 |
| *Reproductive Condition* |  |  |  |  |
| Non-Reproductive | 32.22 | 114.44 | 0.282 | 0.797 |
| Non-Scrotal | -90.33 | 166.74 | -0.542 | 0.63 |
| *Body Mass* | 16.18 | 15.90 | 1.018 | 0.384 |

## Discussion

These results help elicit the importance of understanding sex-specific context in POL research. Although these results suggest considerable variation in FGMs among individuals, animal temperament and FGM concentrations did not show the divergent relationship predicted by alternative reproductive role. Although we did find evidence that reproductive status or age may affect the expression of some behaviours and, ultimately, the syndrome showing the relationship between physiological adaptations and behaviours, we note that there was minimal difference in FGMs between sexes. Given the similarities in basal FGM concentrations between sexes, regardless of reproductive state, we propose that investment in sex-specific POL strategies concerning the traits observed here, may evolve as a result of different directionality of selective pressures. Males may express an increased energetic demand influenced by mate acquisition. In contrast, female conspecifics express a similar rate of energetic stress to accommodate the reproductive costs associated with the care and development of young (Hämäläinen et al. 2018). Although this result does not support the base assumptions of anisogamy and the POLS hypothesis, these findings coincide with previous literature that suggest the emergence of convergent syndromes between sexes (Hämäläinen et al. 2018; Tarka et al. 2018) and elicit the importance of evaluating POL traits under a sex-specific context.

Reproductive costs have been shown to influence the expression of POL traits between sexes (Moshilla et al. 2019; Prabh et al. 2023; Immonen et al. 2018). Generally, males are predicted to express faster strategies because of the energetic costs of dispersal for mate and resource acquisition (Clutton-Brock and Isvaran 2007). Meanwhile, females should express slower strategies and a greater investment in self-maintenance that accommodates the increased energetic costs associated with the development and care of the young (Rogowitz 1996; Moschilla et al. 2019; Réale et al. 2010). Scrotal males and pregnant or lactating females have an increased energetic cost thus, we predicted that individuals in these reproductive stages should have greater FGM levels than non-scrotal and non-reproductive counterparts. Our results show little difference in basal FGM concentrations between individuals during different reproductive stages. Understanding sex-specific ecological context may help further interpret this result. Notably, all individuals compared within this study were surveyed within a breeding season, where all species observed are known to reproduce several times. Thus, the reproductive costs associated with the breeding season may be better informed by examining FGMs and behaviours between distinct breeding and non-breading seasons.

Beyond alternative physiological mechanisms, the POLS also predicts that individual differences in behaviour evolve from trade-offs in energetic demand (Réale et al. 2010; Réale et al. 2007). Males often express a lower level of docility and a higher rate of exploration associated with mate acquisition (Montigilio et al. 2012; Réale et al. 2007). Similarly, females often express a higher level of docility and a lower level of exploration, associated with the protection of young and resulting in greater individual survivorship (Yao et al. 2023; Réale et al. 2007). Given the lack of difference in docility and exploration within the present study, we further suggest that our study species may express a covaried syndrome due to differences in selective pressures influencing reproductive costs. Namely, males may invest more heavily in exploration for mate acquisition; however, because the reproductive output and energetic demand for resource acquisition are consistent between sexes, females may also show similar personalities regarding movement behaviour. Likewise, other environmental pressures may be worth investigating to understand the trade-offs in metabolic stress and behaviour. For example, Stead et al. (2024) found that flying squirrels (*Glaucomys* spp.) experience a peak in fecal cortisol metabolites during autumn, a time that coincides with peak energetic demand associated with food caching rather than reproduction.

Understanding the relationship between environmental pressures and the expression of behavioural and physiological traits is important to understanding how POL traits arise. Indeed, resource quality and availability have been shown to impact trait expression (Bright-Ross et al. 2020; Prabh et al. 2023; Finn et al. 2018). Similarly, the seasonal variation shown in common voles (*Microtus arvalis)* may further correlate with resource availability (Eccard and Herde, 2013). Given that resource availability will fluctuate based on season and undoubtedly depend on population abundance, these are important factors to consider when evaluating POL traits. Indeed, intra- and interspecific competition may result in the expression of various behavioural strategies amongst species, altering movement patterns such that individuals, regardless of sex, must disperse and forage in alternative patterns (Hassel et al. 1994; Wauters et al. 2019). Therefore, resource and habitat availability may further contribute to the expression of similar temperamental traits.

Our results suggested there is a relationship between docility a FGMs, most prominently observed in pregnant red-backed voles and deer mice. We also did not see any significant relationship between FGM levels and the month of sample collection, suggesting low monthly variation during the breeding season. However, further investigation into seasonal variation across winter and summer months, where energetic costs differ, may highlight a more significant relationship (Moffatt et al. 1993). FGM concentrations should increase from spring to summer in most rodents, given the increased energetic demand for reproductive tasks (Stewart et al. 2020; Romero 2002). Veitch et al. (2021) found a significant negative correlation between FGMs and month, attributing the plausible cause as consistent energetic cost in reproduction from May through August. Our results further support this conclusion, as behaviour showed no significant relationship with date in the present study. Similar environmental costs may also explain the lack of sexual differences observed within this study. While females actively investing in the care or development of young may express an increased level of energetic stress (Künkele 2000) similar increased cost may be incurred by males experiencing the associated costs of defense, dispersal, spatial movement, or hormone investment (Millar 1975; Romero 2002).

FGM concentrations and exploration time were positively linked in sub-adult deer mice. This relationship may be explained by immune system maturation (Holt and Jones 2000; Simon et al. 2015) and the necessity of dispersal from individual natal areas (King 1968), further supported by Veitch et al. (2021). Interestingly, we also found a negative relationship between exploration time and FGM concentrations for adults of all three species. While increased dispersal and movement within a home range should increase environmental interactions associated with conflict between conspecifics and potential predators (Mayer et al. 2020), adults should have greater immunocompetence (Webster et al. 2002), and a lower rate of dispersal (King 1968). Therefore, while exploratory behaviours in adults may be greater or equal to younger conspecifics, total FGM production may be comparatively lower.

There was a high degree of variation in FGM concentration and behaviour across individuals, a well-documented phenomenon (Veitch et al. 2021; St. Juliana et al. 2014; Stedman et al. 2017). Indeed, FGM concentration can be influenced by various environmental and genetic factors (Touma and Palme 2005), making conclusions about single observation samples difficult. Likewise, our single season sample size is a limitation when assessing ongoing or re-occurring trends in FGMs and behaviour. Although we did not observe the expected POLS and sex-specific relationships between males and females of the same species, we did see a clear difference in the syndrome among species. While these observations conflict with the predictions of the POLS hypothesis, we note that there was an observable difference in behaviours and glucocorticoid concentrations amongst species. Given the similarities observed between male and female rodents during similar breeding conditions, we suggest 3 considerations for the expression of behaviour and accommodating physiological demand observed within this study. First, a uniformity in selective pressure, such that males and females both experiences increased energetic demand during the breading season may result in convergent behavioural strategies. Second, variation in energetic stress during the breeding season remains consistent, as suggested by Veitch et al. (2021), resulting in the appearance of a uniformity in traits. Finally, although reproduction is an energetically expensive and ecologically important tasks, other or possibly more energetically demanding circumstances may be a larger influence on the expression of POLS traits, as seen in the peak expression of elevated cortisol levels in flying squirrels (Stead et al. 2024). To conclude, although we did not see a definitive adherence to the hypothesis suggested by the POLS framework or anisogamy, we remain enthusiastic about the contextual elements of sex-specific variation in POLS traits and further encourage investigation into ecologically relevant phenomena including a sex-specific lens.

