## Supplementary data C for "Sex differences and behaviour in the pace-of-life of rodents"

**Appendix C: The relationships between total fecal glucocorticoid metabolites and % antibody binding compared to a standard curve generated using Soper, 2021. Shown is the relationship for Deer Mice, Red-Backed Voles and Woodland Jumping Mice for corticosterone or cortisol to validate fecal metabolites.**

Deer Mouse Fecal Corticosterone Parallelism

% Antibody Bound

| log (corticosterone concentration) | Standard curve (y = -17.44ln(x) + 131.22, r² =0.99) | Fecal pool (y = -17.84ln(x) + 117.85, r² = 0.99) |
| --- | --- | --- |
| 7.81 |  | 80.044 |
| 15.63 | 83.869 | 69.599 |
| 31.25 | 70.544 | 57.924 |
| 62.5 | 60.073 | 43.582 |
| 125 | 45.242 | 32.113 |
| 250 | 35.434 | 17.164 |
| 500 | 23.307 | 8.215 |

Red-backed Vole Fecal Corticosterone Parallelism

% Antibody Bound

| log (corticosterone concentration) | Standard curve (y = -17.42ln(x) + 131.81, r² =0.99) | Fecal pool (y = -17.38ln(x) + 119.81, r² = 0.99) |
| --- | --- | --- |
| 7.81 |  | 80.361 |
| 15.63 | 86.577 | 73.352 |
| 31.25 | 70.798 | 61.697 |
| 62.5 | 58.867 | 50.961 |
| 125 | 44.112 | 36.701 |
| 250 | 36.69 | 21.825 |
| 500 | 25.497 | 10.582 |

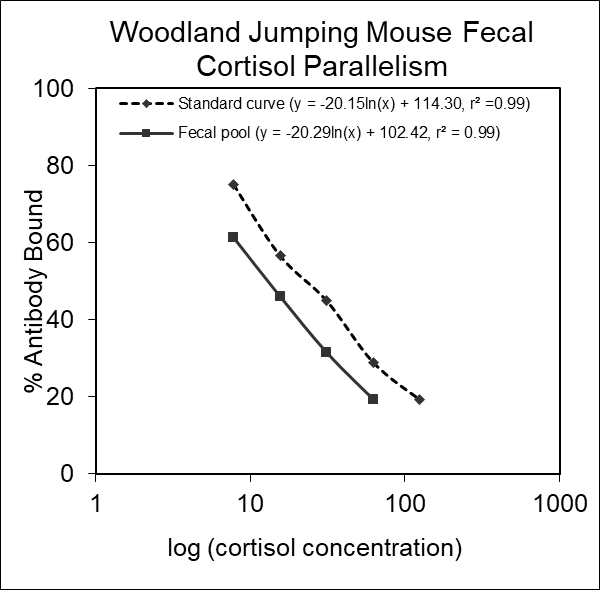

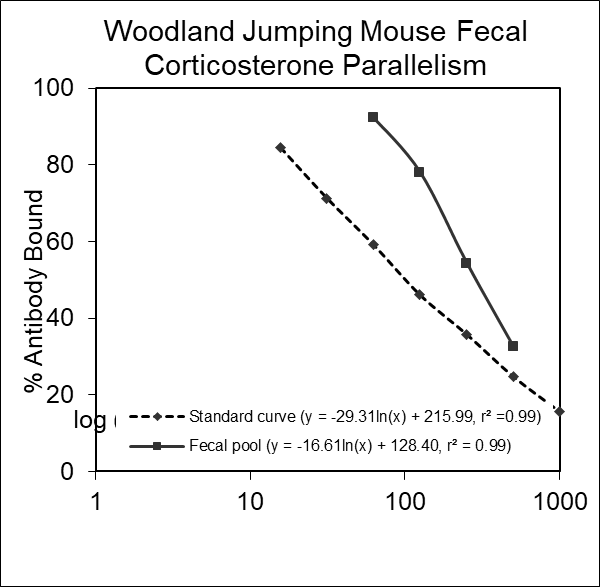

Woodland Jumping Mouse Fecal Cortisol Parallelism

% Antibody Bound

| log (cortisol concentration) | Standard curve (y = -20.15ln(x) + 114.30, r² =0.99) | Fecal pool (y = -20.29ln(x) + 102.42, r² = 0.99) |
| --- | --- | --- |
| 7.81 | 75.143 | 61.42 |
| 15.63 | 56.609 | 46.104 |
| 31.25 | 44.908 | 31.555 |
| 62.5 | 28.881 | 19.388 |
| 125 | 19.172 |  |

Woodland Jumping Mouse Fecal Corticosterone Parallelism

% Antibody Bound

| log (corticosterone concentration) | Standard curve (y = -29.31ln(x) + 215.99, r² =0.99) | Fecal pool (y = -16.61ln(x) + 128.40, r² = 0.99) |
| --- | --- | --- |
| 15.63 | 84.459 |  |
| 31.25 | 71.272 |  |
| 62.5 | 59.158 | 92.324 |
| 125 | 46.085 | 78.073 |
| 250 | 35.859 | 54.37 |
| 500 | 24.886 | 32.505 |
| 1000 | 15.707 |  |
